# Distinct oligodendrocyte populations have spatial preference and injury-specific responses

**DOI:** 10.1101/580985

**Authors:** Elisa M. Floriddia, Shupei Zhang, David van Bruggen, João P. Gonçalves dos Santos, Müge Altinkök, Enric Llorens-Bobadilla, Jonas Frisén, Gonçalo Castelo-Branco

**Affiliations:** Laboratory of Molecular Neurobiology, Department Medical Biochemistry and Biophysics, Karolinska Institutet, Biomedicum, 17177 Stockholm, Sweden; Department of Cell and Molecular Biology, Karolinska Institutet, Biomedicum, 17177 Stockholm, Sweden; Ming Wai Lau Centre for Reparative Medicine, Stockholm node, Karolinska Institutet, 171 77 Stockholm, Sweden

## Abstract

Oligodendrocytes (OLs), the myelinating cells of the central nervous system, are transcriptionally heterogeneous,^1^ the origin and functional consequences of which are unknown. Functional heterogeneity of MOLs might correlate with the local environment or their interactions with different neuron types.^2^ Here, we show that distinct MOL populations have spatial preference in the mammalian central nervous system and differential susceptibility to traumatic spinal cord injury. We also show that the generation of distinct MOL populations is independent of the OPC developmental origin. We found that OPCs originating from the previously described developmental waves^3–5^ have comparable potential to differentiate into the main MOL populations. Furthermore, we found that MOL type 2 (MOL2) is enriched in the spinal cord and almost absent in the brain, while MOL5/6 is enriched with age in all analyzed regions. MOL2 and MOL5/6 also have differential preference for motor and sensory tracts in the spinal cord. In the context of disease, we found that MOL2 and MOL5/6 have differential susceptibility to traumatic spinal cord injury, where MOL2 are lost and MOL5/6 increased their contribution to the OL lineage. Importantly, MOL2 susceptibility is disease specific, as we found MOL2 is not lost in a mouse model of multiple sclerosis. Our results demonstrate that the MOL populations, previously described by single-cell transcriptomics,^1^ have distinct spatial preference and responses. We anticipate our study to pave the way for a better understanding of the MOL populations-specific functional roles in development, health, and disease, allowing for better targeting of the OL subtypes important for the regeneration and repair of the central nervous system.

## Main text

Oligodendrocytes (OLs) metabolically support axons and increase conduction speed.^6^ We have recently reported that the OL lineage is heterogeneous, as it is composed of transcriptionally distinct subpopulations/states during development and disease.^1,7,8^ The heterogeneity of the OL lineage does not lie exclusively within the OL transcriptome. Indeed, developmentally distinct oligodendrocyte progenitor cell (OPC) pools generate OL lineage cells with different abilities to respond to demyelination.^9^ Mature OLs form myelin internodes of various length and thickness, even along the same axon,^10,11^ and show properties regulated at both cell-autonomous and non-autonomous levels.^12,13^

While single-cell RNA-sequencing (scRNAseq) unveiled the transcriptional heterogeneity of the OL lineage and suggested differential enrichment of MOLs in different regions of the CNS,^1^ technical limitations, such as differences in viability of cell subpopulations during tissue dissociation,^14^ might lead to biased cell enrichments. Thus, we assessed the distribution of the OL lineage within white matter (WM) and grey matter (GM) of the brain and spinal cord *in situ* (Extended Data Fig. 1A-B). We performed immunohistochemistry (IHC) and RNAscope in situ hybridization (ISH) for Sox10 as a pan marker of the OL lineage and analyzed confocal images of the corpus callosum (WM), somatosensory cortex (GM) and spinal cord dorsal horn (GM). We analyzed the images with a custom automated pipeline (CellProfiler, Extended Data Fig. 1C-F). Ptprz1 (receptor-type tyrosine-protein phosphatase zeta 1), a marker of OPCs and committed OPCs (COPs),^1,7^ presented a homogeneous distribution across the analyzed regions (Extended Data Fig. 1G, I, K, O. Extended Data Table 1, 2). As expected, we observed a decrease from juvenile to adulthood in the OPCs/COPs (*Ptprz1*^+^-*Sox10*^+^) contribution to the OL lineage cells (*Sox10*^+^), especially in the somatosensory cortex (Extended Data Fig. 1G, I, K, O. Extended Data Table 1, 2). Among the six transcriptionally distinct mature oligodendrocyte populations previously described, MOL1, MOL2, and MOL5/6 present the most distinct gene marker modules,^1,7,8^ therefore we analyzed their spatial distribution. *Egr2* (*Early Growth Response 2*; also known as *Krox20*) is expressed specifically by MOL1 in the OL lineage^1^ and we observed an even distribution of MOL1 within the analyzed regions (Extended Data Fig. 1H, J, L, P. Extended Data Table 1, 2). *Klk6* (*Kallikrein Related Peptidase 6*) is a distinct marker for MOL2.^1,7,8,15^ Klk6 has been previously associated with demyelinating pathology in experimental autoimmune encephalomyelitis (EAE) and spinal cord injury (SCI).^16,17^ Strikingly, we observed that *Klk6*^+^ MOL2 is a population specifically enriched in the spinal cord and almost absent in the brain (Fig. 1A-B, Extended Data Fig. 1M, Q. Extended Data Table 1, 2). In contrast, MOL5/6, expressing *Ptgds* (*Prostaglandin D2 Synthase*),^1,7,8^ showed a dynamic contribution to the OL lineage, increasing along time and following the myelination temporal pattern. Indeed, at juvenile (P20), MOL5/6 are more abundant in the spinal cord, where myelination is visible early on after birth (P5-6), compared to the brain, where myelination is visible around P10-15. In adulthood (P60), MOL5/6 is the main population contributing to the OL lineage in both brain and spinal cord, being most abundant in the corpus callosum (Fig. 1C-D, Extended Data Fig. 1N, R. Extended Data Table 1, 2). Thus, MOL2 and MOL5/6 present distinct distributions and spatial preference across the CNS, contrary to MOL1.

**Fig. 1.**
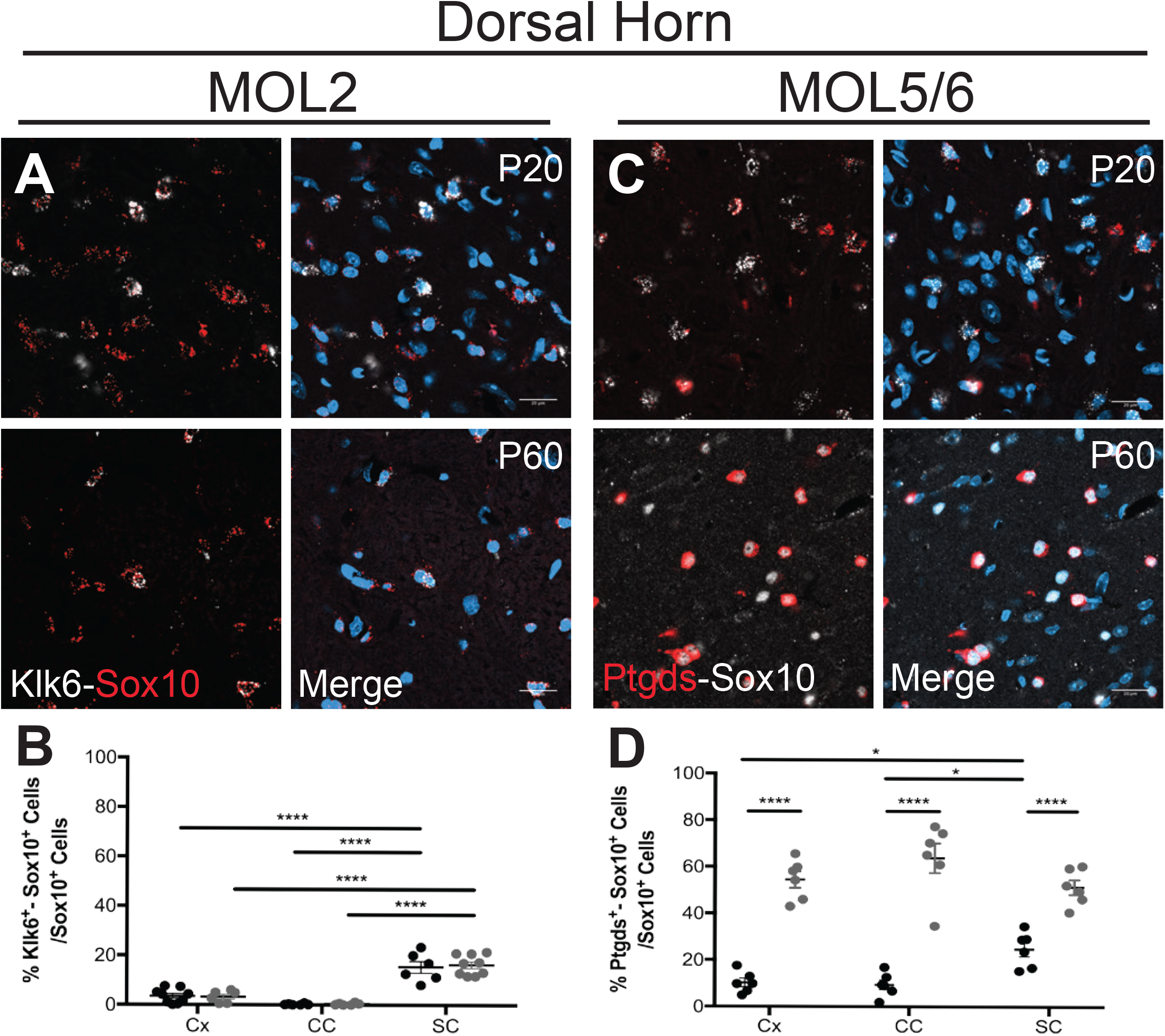
Specific mature OL populations have spatial preference in the juvenile and adult central nervous system. (**A, C**) Confocal representative images of the distribution of MOL2 (Klk6^+^ OL lineage cells. A), and MOL5/6 (Ptgds^+^ OL lineage cells. C) in the dorsal horn (grey matter) of the juvenile (P20) and adult (P60) spinal cord. Scale bar = 20 μm. (**B, D**) Quantification of the MOL2 (B), MOL5/6 (D) distribution in the cortex, corpus callosum, and dorsal horn in juvenile (P20) and adulthood (P60). Percentage of the population is calculated on the total number of OL lineage cells (Sox10^+^ cells) in the analyzed region. Data are presented as Mean ± SEM. n = 6-9 animals per condition. Asterisks indicate a significant difference between conditions (*p ≤ 0.05, ****p ≤ 0.0001, 2-way ANOVA with Sidak’s correction). Cx = cortex, CC = corpus callosum, SC = spinal cord.

The OL lineage derives from distinct progenitor domains and developmental stages,^3^ which might have a role in the specification of OPCs into the MOL populations and therefore a role in the observed spatial preference. To investigate this possibility, we first isolated the OL lineage from Emx1::Cre-Sox10::GFP-TdTom mice,^1,5,7^ and performed scRNAseq on ventrally (eGFP^+^) and dorsally (TdTom^+^) derived OL lineage cells. We analyzed the P60 corpus callosum (Fig. 2A) since this region has representation of OLs derived from both the cortical plate (dorsal domain) and the lateral and medial ganglionic eminences (ventral domains of the embryonic forebrain).^5^ As expected,^5^ a greater proportion of TdTom+ OL lineage was obtained, (Fig. 2A, D. Extended Data Fig. 2B, Extended Data Table 3). Clustering analysis with GeneFocus^7,18^ of 2,853 OL lineage cells led to the identification of the previously identified OL lineage subpopulations (Fig. 2B, Extended Data Fig. 2A-B).^1,19^ Importantly, the contribution of ventrally- and dorsally-derived OL lineage cells to each cluster was comparable (Fig. 2C-D), suggesting that the developmental waves have similar potential to generate the transcriptionally distinct OL lineage subpopulations.

**Fig. 2.**
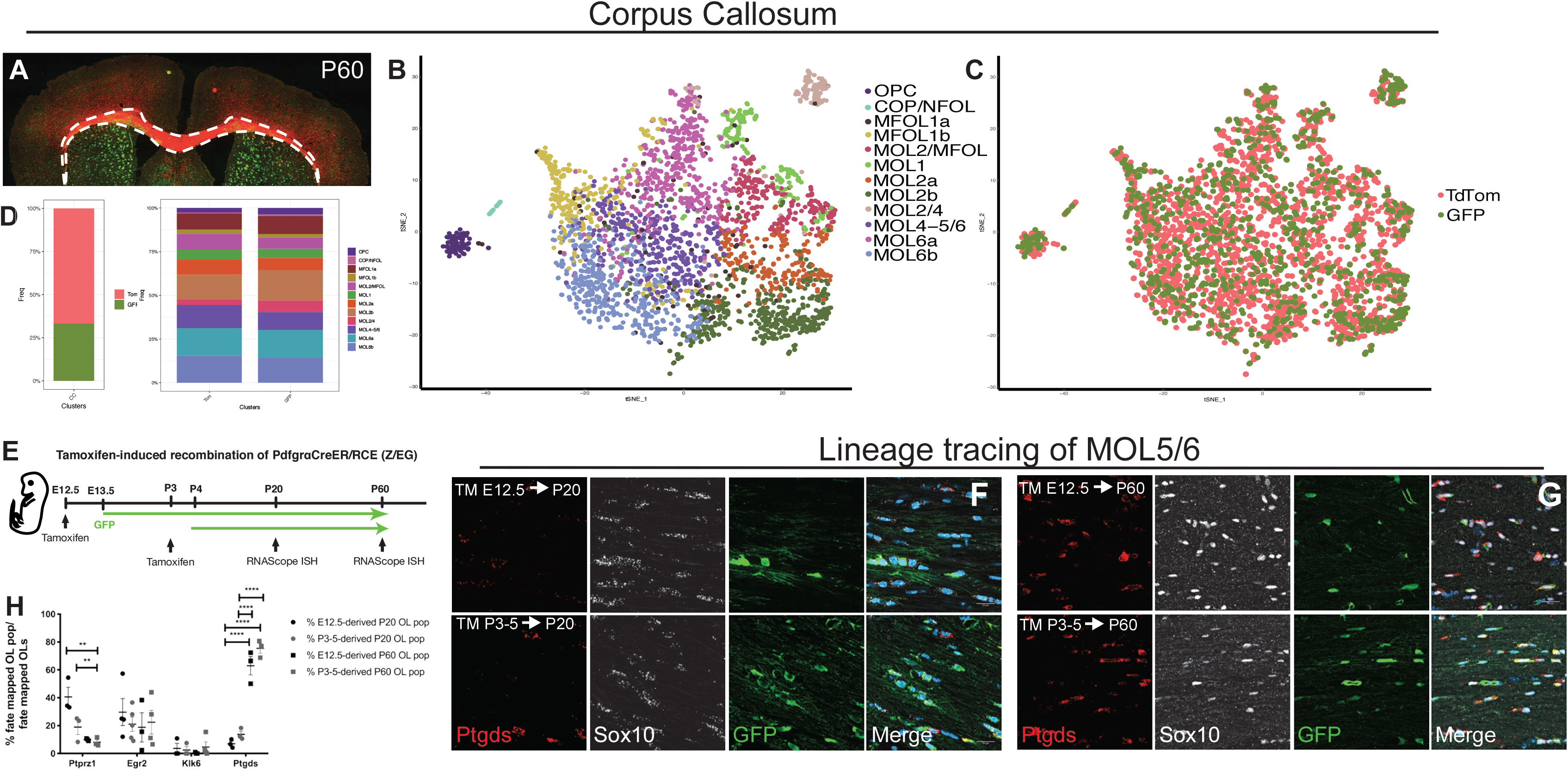
The developmental origin does not specify OPCs into distinct MOL subpopulations. (**A**) Confocal representative images of the P60 brain from Emx1::Cre-SOX10::GFP-TdTom mouse. Dashed outline highlights the corpus callosum and dissected region used for scRNAseq. (**B-C**) t-Distributed stochastic neighbor embedding (t-SNE) plots showing the OL lineage composition determined by hierarchical clustering (B) and the TdTom^+^ and GFP^+^ OL lineage cells contribution to the clusters (C). (**D**) Frequency distribution of the TdTom^+^ and GFP^+^ OL lineage cells forming the major clusters. (**E**) Schematic overview of the fate mapping experimental design. Green arrows shows the GFP expression timeline 24h delayed from the time of tamoxifen injection. (**F-G**) Confocal representative images show the MOL5/6 (Ptgds^+^-GFP^+^ OL lineage cells differentiated from pre- (TM E12.5) or postnatal (TM P3-5) OPCs. Scale bar = 20 μm. Sox10 was detected by RNAscope ISH in F and by IHC in G. (**H**) Percentages of the fate mapped OPCs-COPs (Ptprz1^+^-GFP^+^ OL lineage cells), MOL1 (Egr2^+^-GFP^+^ OL lineage cells), MOL2 (Klk6^+^-GFP^+^ OL lineage cells), and MOL5/6 (Ptgds^+^-GFP^+^ OL lineage cells) populations are calculated on the total number of fate mapped OL lineage cells (Sox10^+^-GFP^+^ cells) in the juvenile and adult corpus callosum. Data are presented as Mean ± SEM. n = 3-5 animals per condition. Asterisks indicate a significant difference between conditions (**p ≤ 0.01, ***p ≤ 0.0001, 2-way ANOVA with Sidak’s correction). TM = tamoxifen, OPC = oligodendrocyte progenitor cell, COP = committed OPC, NFOL = newly formed oligodendrocyte, MFOL = myelin forming oligodendrocyte, MOL = mature oligodendrocyte, TdTom = tandem duplicated tomato, GFP = green fluorescent protein.

Ventrally-derived and dorsally-derived OL lineage cells arise during embryonic and early-postnatal development, respectively.^3,4,20^ We thus further performed lineage tracing using the Pdgfrα::CreERT1-loxP-GFP mouse model^21^ to fate map the progeny of pre-OPCs appearing at E12.5^7^ or OPCs at P3-5 (Fig. 2E). Postnatal recombination of the Pdgfrα::CreER^T1^-GFP mice results in the assessment of OPCs arising at both postnatal stages and remaining pre-OPCs, while E12.5 recombination allows the assessment of ventral pre-OPCs arising exclusively from this embryonic stage. Our recombination strategy labelled the majority of the lineage in the juvenile CNS (95.43 ± 6.22% of the Sox10+ cells were positive for the eGFP reporter in the cortex, 62.24 ± 2.35% in corpus callosum, and 96.05 ± 1.15% spinal cord, Extended Data Fig. 3A). However, we observed sparser labeling in adulthood (62.38 ± 19.9% of the Sox10^+^ cells were also positive for the eGFP reporter in the cortex, 44.16 ± 9.04% in the corpus callosum, and 24.32 ± 8.26% in the spinal cord. Extended Data Fig. 3B). This differential labeling suggests a considerable contribution of non-recombined OPCs (GFP^−^) within the process of life-long continuous addition of new myelinating OLs for myelin homeostasis and in response to experience.^22,23^ Importantly, we observed that a subsets of E12.5 pre-OPCs and their progeny do not disappear later in life in any of the analyzed regions (Fig. 2F-H, Extended Data Fig. 3), as previously suggested.^3–5^

We did not observe significant difference in the contribution of the two developmental waves to the MOL1, MOL2, and MOL5/6 populations (Fig. 2H, Extended Data Fig. 3C-Q). While most of the MOL2 in the dorsal spinal cord were derived from E12.5 pre-OPCs at P20, the contribution from postnatally fate mapped OPCs reached comparable levels in adulthood (19.09 ± 4.95% and 23.35 ± 1.28% from the postnatally- and embryonically-derived MOL2, respectively. Extended Data Fig. 3M, O-P). This suggests that postnatally-derived OPCs start differentiating into MOL2 later than embryonically-derived pre-OPCs. In sum, our scRNAseq and lineage tracing data suggest that the domain and time of origin does not influence the OPC specification towards distinct MOL populations. Rather, exposure of OPCs to extrinsic cues during critical windows of migration or differentiation might contribute to MOL diversification at their final localization.

Given the preferential localization of MOL2 in the spinal cord (Fig. 1A-B), we further investigated whether MOL2 and MOL5/6 spatial preference relates to functionally distinct regions and tracts in the spinal cord (Fig. 3A). Strikingly, we observed an enrichment of MOL2 in the WM compared to the GM of the spinal cord (Fig. 3B, C. Extended Data Table 4), maintained with age (Fig. 3C. Extended Data Table 4). On the contrary, the MOL5/6 population is enriched in the GM compared to the WM of the spinal cord (Fig. 3F, G. Extended Data Table 4). Interestingly, the MOL2 population decreases in the GM of the spinal cord during adulthood (Fig. 3C. Extended Data Table 4). In accordance with previous reports,^24–28^ in the dorsal white matter, the OL lineage density around the dorsal columns and dorsal corticospinal tract corresponds to the development of myelination (Extended Data Fig. 1F). In the descending dorsal motor corticospinal tract, we observed an increase in the OL lineage with age (Extended Data Fig. 1F. Extended Data Table 4) and constant distribution in the ascending sensory dorsal columns (Extended Data Fig. 1F. Extended Data Table 4). Importantly, we observed they have adjacent distribution for MOL2 and MOL5/6, as when we compared their spatial preference for the GM and WM of the spinal cord (Fig. 3B-C, F-G). Indeed, MOL2 are preferentially found in the dorsal columns (Fig. 3D, E), while MOL5/6 preferentially localize in the dorsal corticospinal tract (Fig. 3H, I. Extended Data Table 4).

**Fig. 3.**
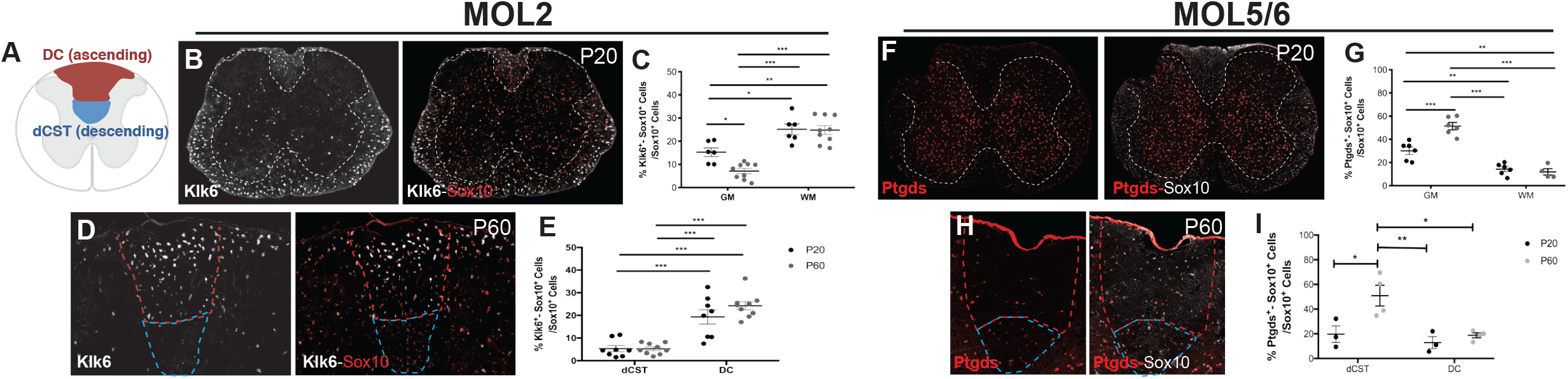
MOL2 and MOL5/6 are specifically enriched in adjacent regions of the juvenile and adult spinal cord. (**A**) Schematics of a coronal section of the spinal cord. Highlighted in blue and red the white matter regions where the axons forming the dorsal columns and dorsal corticospinal tract run, respectively. (**B, D**) Confocal representative images show the enrichment of MOL2 (Klk6^+^ OL lineage cells) in the white matter of the spinal cord (B) and at the level of the dorsal columns (D). (**F, H**) Confocal representative images show the enrichment of MOL5/6 (Ptgds^+^ OL lineage cells) in the grey matter of the spinal cord (F) and at the level of the dorsal corticospinal tract (H). Scale bar = 100 μm. (**C, E, G, I**) Quantification of the MOL2 and MOL5/6 distribution in the white and grey matter of the spinal cord (C, G) and at the level of the dorsal corticospinal tract and dorsal columns (E, I) in juvenile (P20) and adulthood (P60). Percentage of the population is calculated on the total number of OL lineage cells (Sox10^+^ cells) in the analyzed region. Data are presented as Mean ± SEM. n = 3-9 animals per condition. Asterisks indicate a significant difference between conditions (*p ≤ 0.05, **p ≤ 0.01, ***p ≤ 0.001, 2-way ANOVA with Sidak’s correction). GM = grey matter, WM = white matter, DC = dorsal columns, dCST = dorsal corticospinal tract.

The clear spatial preference of MOL2 and MOL5/6 for different regions and tracts of the spinal cord together with the importance of extrinsic, rather than intrinsic, factors for their diversification suggest that the local environment might have an important role in the physiological properties of MOLs. This might also lead to population specific responses in disease. Traumatic injury of the spinal cord is a chronic pathological condition that leads to loss of locomotor and sensory functions due largely to Wallerian degeneration.^29^ We took advantage of two models of traumatic injury: dorsal funiculi transection (an axotomy model of mild severity. Fig. 4A) and severe contusion injury (Fig. 4H). During the remyelination stage following injury, we observed that the OL lineage is still well represented distal to and at the injury site (Fig. 4D-E, K).^30–34^ However, in both injury models, we observed a specific loss of MOL2 at the injury site (Fig. 4C-B, F-G,I-J, L), maintained along time (Fig. 4F-G). On the contrary, MOL5/6 reached a higher contribution to the OL lineage at the injury site (Fig. 4C, F-G, J, L) compared to the intact spinal cord (Fig. 3F-I), despite the absence of intact axons. We also assessed the spatial distribution of MOL2 and MOL5/6 in regions of Wallerian degeneration rostral and caudal to the injury site (Fig. 4C, F-G; Extended Data Fig. 4). We observed that the spatial preference of MOL2 and MOL5/6 for the dorsal columns and dorsal corticospinal tract was unaffected by the injury (Extended Data Fig. 3). The contribution of MOL5/6 in the dorsal funiculi rostral and caudal to the injury did not change following injury (18.8 ± 2.82% and 18.75 ± 2.9% at 3 and 5 mpi, respectively. Fig. 4F-G and Extended Data Fig. 3), and remained similar to their contribution in the white matter of the intact adult spinal cord (Fig. 3F-G. Extended Data Table 4). Surprisingly, the contribution of MOL2 to the OL lineage in the same regions increased with time (Extended Data Fig. 4B-E). Indeed, 5 mpi, the MOL2 cell density is even greater than found in the WM of the intact adult spinal cord (Fig. 3B-C. Extended Data Table 4).

**Fig. 4.**
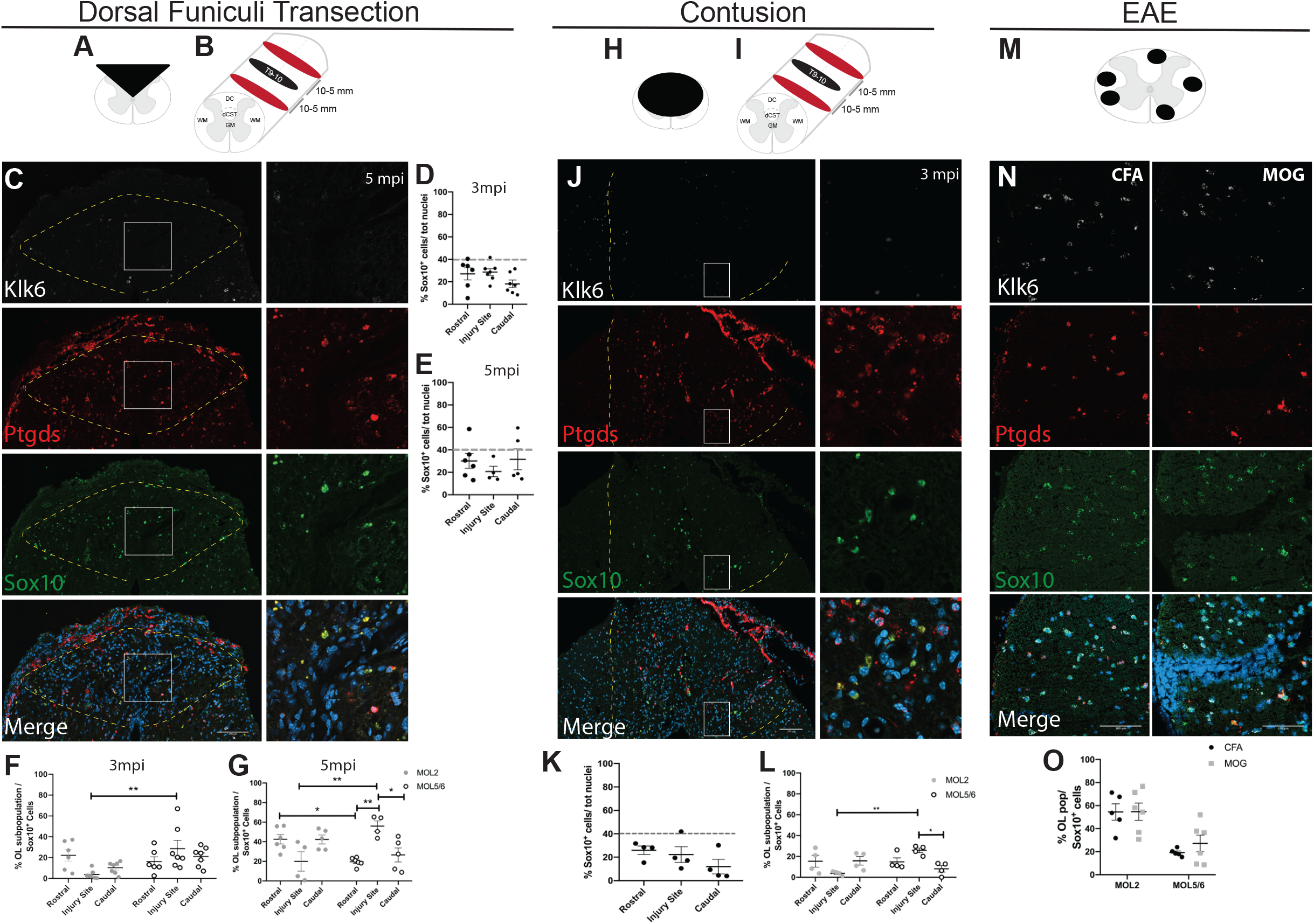
MOL2 and MOL5/6 show differential and disease-specific susceptibility. (**A-B, H-I, M**) Schematic of the analysis lesion model (A, H, M), level of the lesions (T10. B-I) and distance of the rostral and caudal analyzed segments (B-I). (**C, J, N**) Confocal representative images of the lesions following dorsal funiculi transection (C) and contusion (J) injuries, as well as in EAE (N), showing the specific loss of MOL2 (Klk6^+^ OL lineage cells) and the high repopulation by MOL5/6 (Ptgds^+^ OL lineage cells) of the lesions following traumatic injury of the spinal cord. Yellow dashed lines highlight the lesion sites. White rectangles highlight the regions shown in higher magnification. Scale bar =100 μm. (**D-G, K-L, O**) Quantification of the OL lineage (D-E, K) and the MOL subpopulations away and at the traumatic injury sites (F-G, L) and in EAE lesions (O). Percentage of the MOL2 and MOL5/6 subpopulations was calculated on the total number of OL lineage cells (Sox10^+^ cells). Dashed grey line marks the average percentage of the OL lineage cells in the intact adult spinal cord (D-E, K). Data are presented as Mean ± SEM. n = 4-7 animals per condition. Asterisks indicate a significant difference between conditions (*p ≤ 0.05, **p ≤ 0.01, 2-way ANOVA with Sidak’s correction). SCI = spinal cord injury, mpi = months post-injury, T10 = thoracic vertebra 10, CFA = Complete Freund’s Adjuvant, MOG35-55 = Myelin Oligodendrocyte Glycoprotein, peptide fragment 35-55, PTX = Pertussin Toxin.

To assess whether the MOL2 loss is disease specific or a general signature of demyelinating and axonal pathology, we took advantage of experimental autoimmune encephalomyelitis (EAE). EAE is a mouse model of CNS inflammation presenting multifocal lesions characterized by extensive oligodendrocyte loss and Wallerian degeneration. Unlike in the spinal cord injury model, we observed a similar distribution of MOL2 and MOL5/6 in the intact or lesioned white matter of the spinal cord in mice immunized with CFA or MOG35-55, respectively (Fig. 4N-O), consistently with our previous transcriptomic analysis identifying a MOL2 population enriched in the spinal cord of EAE mice.^18^ In summary, our data indicates that MOL2 is specifically lost and increase their contribution to the OL lineage in regions of Wallerian degeneration following traumatic spinal cord injury. MOL2 might be associated with circuit remodeling, resulting from sprouting of intact axons proximal to the injury site, while the injury site might present factors stimulating MOL5/6 survival or resident OPCs preferentially differentiation to MOL5/6.^12,35,36^ Altogether, our data show OL lineage populations present disease specific susceptibility.

Here, we unveiled that distinct MOL populations have spatial preference in the mammalian central nervous system and differential susceptibility to traumatic spinal cord injury. Additionally, we show that the generation of distinct MOL populations is independent of the OPC developmental origin, and the MOL population-specific susceptibility to traumatic injury. Our data suggest that MOL5/6 might be an OL population associated with adaptive myelination, as MOL5/6 increases in the corpus callosum and CST from juvenile to adulthood. Indeed, the tracts forming both the corpus callosum and the CST maintain plasticity throughout life, essential for acquisition and maintenance of new motor skills with continuous OL turnover, suggesting that myelin needs to be constantly remodeled (adaptive myelination) in those regions.^22,27,28^ In contrast, we show that MOL2 preferentially locate in the dorsal columns (sensory tracts), suggesting that MOL2 might be associated with more stable neural circuits or production of myelin favoring fast conduction. Indeed, proprioceptive and mechanoreceptive long projecting fast axons are myelinated early in development and reach complete myelination by juvenile,^28,37^ correlating with the high and stable enrichment of MOL2 in this region. Additionally, MOL2 are transiently over-produced in the grey matter of the spinal cord, suggesting that MOL2 selection and correct localization might be regulated by intrinsic mechanisms, such as programmed cell death.^38,39^ This observation is in line with the lack of developmental OPC specification into distinct mature OL populations. Our observations fit a model where the mechanisms regulating differentiation of the OL lineage are independent on the domain of origin,^3,4,20^ differently than it is for neurons.^40,41^

The proportion of MOL2 decreases after traumatic spinal cord injury but not in EAE. While both pathologies have common features, such as axonal and myelin loss, they also present differences which might contribute to MOL2s differential susceptibility. So-called quiescent oligodendrocytes has been previously described in Wallerian degeneration.^42–44^ In accordance, we observed a considerable presence of MOL2 and MOL5/6 in regions where the dorsal columns and corticospinal tracts undergo axonal degeneration. Thus, Wallerian degeneration most likely does not affect the relative contribution of MOL2 and MOL5/6 to the OL lineage.

Our results show that specific mature OL populations present differential response to traumatic injury. MOL2 and MOL5/6 might have differential contribution to remyelination, axonal support, action potential conduction, and synapse formation inhibition, processes with potential high impact for regenerative medicine. We anticipate our study to pave the way for further understanding of the MOL populations-specific functional roles in development, health, and disease, allowing for better targeting of the OL subtypes important for the regeneration and repair of the central nervous system.

## Acknowledgements

We would like to thank Sarah Foerster, Richa B. Tripathi, William D. Richardson, and Robin J.M. Franklin for the fruitful discussions and sharing the Emx1::Cre-Sox10::Cre-LoxP-GFP-STOP-TdTom line, Rashid Holtinkoski, Eneritz Agirre, Ana Mendanha Falcão, Alessandra Nanni, Johnny Söderlund, Ahmad Moshref, Tony Jimenez-Baristain, the Facility Management and Administration at Biomedicum (Karolinska Institutet, Stockholm) for laboratory management and support. Dr. Göran Månsson and Connla Edwards at the Biomedicum Imaging Core Facility (Karolinska Institutet, Stockholm), Dr. Jaromir Mikes at the Mass Cytometry National Facility (SciLife Lab, Stockholm); Drs. Karolina Wallenborg, Anna Juréus, Marcela Ferella at the Eukariotic Single Cell Genomics facility (SciLife Lab, Stockholm); Katarina Ericsson, Kristoffer Tenebro Berglund, Johanna Hornstrand at the Comparative Medicine Biomedicum facility (Karolinska Institutet, Stockholm) and their respective facility managements, the National Genomics Infrastructure and Uppmax for providing assistance in massive parallel sequencing and computational infrastructure.

The bioinformatics computations were performed on resources provided by the Swedish National Infrastructure for Computing at UPPMAX, Uppsala University. Work in G.C.-B.’s research group was supported by Swedish Research Council (grant 2015-03558), European Union (Horizon 2020 Research and Innovation Programme/ European Research Council Consolidator Grant EPIScOPE, grant agreement number 681893), Swedish Brain Foundation (FO2017-0075), Ming Wai Lau Centre for Reparative Medicine, Strategic Research Programme in Neuroscience (StratNeuro) and Karolinska Institutet.

## Author Contributions

E.M.F. and G.C.-B. conceived the project, designed the study and interpreted results. E.M.F. and S.Z. performed experiments and analyzed data. J.P.G.S., M.A and E.L.B. performed experiments. D.vB. analyzed data. E.M.F., S.Z., D.vB., E.L.B., J.F., and G.C.-B. discussed the results of the study. E.M.F. and G.C.-B. wrote the manuscript with feedback from all co-authors.

## Materials and Methods

### Animals

All experimental procedures were performed following the guidelines and recommendation of local animal protection legislation and were approved by the local committee for ethical experiments on laboratory animals in Sweden (Stockholms Norra Djurförsöksetiska nämnd).

Mouse lines used in this study are Pdgfrα::CreER^T1^-RCE, Pdgfrα::CreER^T1^-ROSA26::LoxP-GFP,^21^ Sox10::CreER^T2^-ROSA26::LoxP-GFP,^45,46^ available at The Jackson Laboratory, and Emx1::Cre-Sox10::Cre-LoxP-GFP-STOP-TdTom.^5^ Mice were used with the Cre allele in hemizygousity and the reporter gene allele in either hemizygousity or homozygousity.

Animals were sacrificed at juvenile (P20-21) and adult stages (P60), injuries were performed on adult mice (P60-P150) and sacrificed three- or five-months post-injury, EAE was induced in adult mice (P90) and terminated when the MOG35-55-immunized mice reached a clinical score of 3.0 (limp tail and complete paralysis of the hind legs. P100-110). Both sexes were included in the experiments, except for the contusion spinal cord injury paradigm, where only female mice were used. The following light/dark cycle was used: dawn 6:00-7:00, daylight 7:00-18:00, dusk 18:00-19:00, night 19:00-6:00. Mice were housed to a maximum number of 5 per cage in individually ventilated cages with access to food and water ad libitum.

### Lineage tracing

To fate map the OPCs generated at E13.5 and present at P3-5, Pdgfrα::CreER^T1^-RCE, Pdgfrα::CreER^T1^-ROSA26::LoxP-GFP mice were used.^21^ Time-mated females were injected i.p. with 1 mg of tamoxifen (Sigma) at pregnancy day E12.5 or 2 mg once daily when pups were P3 to P5 (tamoxifen reaching the pups via the mother’s milk). At this low dose (1mg) of tamoxifen during pregnancy (equivalent to 33 mg/kg of body weight) was used to restrict the labeling to the first appearing OPCs. Indeed at this low dose, tamoxifen is metabolized within 24-36 h after injections (therefore E13.5-E14.0.^47,48^). The litters were then sacrificed as juvenile (P20) or young adults (P60) and brains and spinal cords collected for tissue analysis by RNAscope ISH coupled with IHC.

### Spinal cord injury and post-operative care

#### Dorsal funiculi transection

Mice were kept under anesthesia with a 2% isoflurane/O2 mixture at 1 litre/minute, after two-three minutes induction with 5% isoflurane/O2 mixture at 1 litre/minute. Body temperature was maintained by keeping the mice on a heating pad (37-39°C) during the whole procedure. The injury site was mid-thoracic (T10). The fur was shaved and the area disinfected with Clinical (DAX. twice) and 70% EtOH (once). The skin was incised, the superficial fat displaced, the prominent vessel (between T7 and T8) was identified and used as reference point. Then, the muscle tissue over T9-11 removed to expose the vertebrae. A T10 laminectomy was performed, the dura mater was removed, and the dorsal funiculi transection was performed with a microknife. Dorsal funiculi transection damages dorsal columns, dorsal corticospinal tract, and partially the dorsal horns. After surgery, the mice were injected i.p. with Buprenorphine (Temgesic^®^) 0.01 mg/kg of body weight and s.c. with 0.5 ml of 0.9% saline solution. The mice were then placed in their home cages and monitored until fully recovered from anaesthesia. During their postoperative care, mice underwent daily checks for general health status, weight loss, mobility, wounds, swelling, infections, or autophagy of the toes. When mice lost weight after surgery, their diet was supplemented with DietGel^®^ Recovery (Clear H_2_O). The mice used in this study did not show self-induced wounds or autophagy of the toes or wound infections.

#### Contusion

Mice were deeply anesthetized with isoflurane and provided with pre-operative analgesia (Buprenorphine, Schering-Plough, 0.1 mg/kg body weight and Carprofen, Pfizer, 5 mg/kg body weight). A laminectomy was performed at the T8-T9 level to expose the dorsal portion of the spinal cord. the vertebral column was then stabilized with clamps (Precision System Instrumentation, PSI) placed on the foramen of T8 and T10. After the administration of local anaesthesia (Xylocaine/Lidocaine, AstraZeneca, 10 mg/ml, 2 drops on the spinal cord surface), a 70 kDyne contusion was delivered medially over T9 using the Infinite Horizon Impactor (IH-0400, PSI) equipped with a 1.3mm tip. The wounds were then sutured and mice placed in a heated cage until they regained consciousness after which they were transferred to their home cage that was equipped with an elevated floor grid. During the first three days of post-operative care, animals received additional hydration (500ul saline injection daily), antibiotic treatment (Sulphadizine/Trimethoprim, Tribrissen vet., MSD, 100mg/kg body weight per day) and analgesia (Buprenorphine, Schering-Plough, 0.1 mg/kg body weight and Carprofen, Pfizer, 5 mg/kg body weight per day). Bladders were expressed three times per day during the first three days and twice per day until they regained bladder control. Their diet was supplemented with a high energy nutritional supplement (DietGel Boost, Clear H2O) during the first week and starting from 1-2 days prior to the surgery. Body weight was monitored daily during the first week and weekly thereafter. Animals that lost more than 15% of their pre-operative body-weight were euthanized and not included in the study.

### Experimental Autoimmune Encephalomyelitis

For the induction of chronic EAE, animals were injected subcutaneously with an emulsion of MOG35-55 in CFA or vehicle alone (EK-2110 kit, Hooke Laboratories), followed by the i.p. administration of pertussin toxin in PBS (0.2 μg per animal) for two consecutive days (accordingly to manufacture’s instructions). Animals were checked daily to assess general health status, weight loss, wounds, swelling, or infections. When mice lost weight after surgery, their diet was supplemented with DietGel^®^ Recovery (Clear H2O). One mouse showed signs of infection at the subcutaneous injection site and was humanly terminated. The mice were also daily score for clinical sign of EAE (accordingly to manufacturer’s instructions).

### Tissue preparation and sectioning

At the end of the experiments for the visualization in tissue of the OL lineage populations, the animals were deeply anesthetized with ketamine (120 mg/kg of body weight) and xylazine (14 mg/kg of body weight) and transcardially perfused with 0.1 M PBS followed by 4% PFA in PBS (pH 7.4 for both solutions). Brains and spinal cords were dissected and post-fixed in 4% PFA in PBS (pH 7.4) at 4°C overnight and cryoprotected in 30% sucrose for 48-36 hours. Tissue was embedded in Tissue-Tek^®^ O.C.T. compound (Sakura). Both brains and spinal cords were coronally cryosectioned (20 and 16 μm, respectively) in 1:10 series. Sections were stored at −80°C until further use.

### Tissue dissociation and single-cell RNAseq

The corpus callosum of P60 Emx1::Cre-Sox10::Cre-LoxP-GFP-STOP-TdTom^5^ was microdissected from two mice and the tissue dissociated into a single-cell suspension, as previously described.^1^ Briefly, mice were transcardially perfused with ice-cold oxygenated artificial cerebrospinal fluid (22 mM NaCl, 0.63 mM KCl, 0.4 mM NaH_2_PO_4_ * 2H_2_O, 6.5 mM NaHCO_3_, 25 mM Saccharose, 5 mM Glucose, 0.5 mM CaCl_2_, 4 mM MgSO_4_. pH 7.3) and the brains collected. The tissue was sectioned at the vibrotome in ice-cold artificial cerebrospinal fluid and the corpus callosum microdissected. Tissue dissociation was performed with the Adult Brain Dissociation Kit (Miltenyi Biotec) following manufacturer’s instructions (red blood cells removal step was not included). After debris removal, the cells were resuspended in ice-cold 1% BSA in artificial cerebrospinal fluid, filtered with 30 μm filter (Sysmex Partec) and FACS sorted with the BD Influx System (USB. BD FACS™) to separate GFP^+^ and TdTom^+^ OL lineage cells.

The sorted cells were processed with the Chromium Single Cell A Chip kit v2 and library prep with the Chromium Single Cell 3’Library & Gel Beads kit v2 (10X Genomics) accordingly to manufacturer’s instructions. A total of 3,000 cells for each sample was loaded on the Chromium Single Cell A Chip, although a lower number of cells was recovered in singlet and passed the quality control. The scRNAseq dataset is available at GEO (NCBI). GSE128525^1^.

### RNAscope in situ hybridization (ISH) and immunostaining (IHC)

RNAscope ISH was performed using the RNAscope^®^ Multiplex Fluorescent Detection Reagents Kit v2 (ACD Biotechne) on PFA fixed juvenile, adult, and injured brains and spinal cords accordingly to manufacturer’s instructions with some modifications. Briefly, after treatment with boiling 1X target retrieval, the sections were incubated with Protease IV for 20 min at RT, followed by washing and the indicated hybridization and amplification steps. Probes used in this study were designed for mouse Sox10-C1 or -C2 (ACD Biotechne, 435931), Ptgds-C1 (ACD Biotechne, 492781), Klk6-C3 (ACD Biotechne, 493751), Egr2-C3 (ACD Biotechne, 407871), Ptprz1-C1 (ACD Biotechne, 460991).

For lineage tracing experiments and the identification of P20 and P60 OL lineage within tissue from the Pdgfrα::CreERT1-RCE::LoxP-GFP mice,^21,49^ the RNAscope ISH was coupled with IHC to detect the GFP reporter or endogenous Sox10. Briefly, after hybridization and amplification steps to detect the mRNA of the target gene markers, the sections were blocked in 5% normal donkey serum (NDS), 0.03% Triton X100 in PBS for 1h at RT and incubated with chicken anti-GFP (AbCam, ab 13970) or goat anti-Sox10 (Santa Cruz, sc-17342) primary antibodies 1:200 in 2% NDS, 0.03% Triton X100 in PBS, O.N. at RT. The following day, the sections were incubated with goat anti-chicken AlexaFluor 488 conjugated (AbCam, ab150169) or donkey anti-goat AlexaFluor 647 conjugated (LifeTech, A21447) secondary antibodies 1:500 in 2% NDS, 0.03% Triton X100 in PBS, 1h at RT and counterstained with DAPI (1:5,000 in PBS) for 2 min. IHC washing steps were performed with 0.05% Tween-20 in PBS.

### Image acquisition

Fluorescent images were acquired using the LSM800 confocal microscope set up (Zeiss). To obtain an optimal balance between resolution of RNAscope signal and imaged area, tiled images were acquired with a 40X water objective. The z-stack was kept to 2-3 focal planes with 40 μm step to reduce the probability of false positive cells after image maximum projection.

### Image analysis

Confocal images were processed with FiJi (ImageJ, NIH) to select the regions of interest (ROIs). ROI images were segmented with a customized CellProfiler pipeline. Briefly, the signals from the individual channels (DAPI, markers, GFP) were segmented; OL lineage cells (Sox10^+^) were identified using the masking option with the DAPI counterstain. Then, the specific OL lineage populations were identified using the relate objects options with the Sox10^+^ or Sox10^+^-GFP^+^ cells. Overlay images of the identified objects were exported and used to assess the percentage of cell segmentation error. Spreadsheets containing the number of parent cells (DAPI, Sox10^+^ or Sox10^+^-GFP^+^ cells) and child objects (Ptgds^+^, Klk6^+^, Egr2^+^, Ptprz1^+^) were exported and used to calculate the percentage of each population. Based on the average gene expression in scRNAseq dataset^1^ we used a cut off of 12, 4, 3, and 7 molecules of Ptgds^+^, Klk6^+^, Egr2^+^, Ptprz1^+^, respectively, per cells to call the analyzed OL lineage populations.

We manually assessed the percentage of error for the automated cell segmentation and attribution on 18 representative images (six images per analyzed region). We recorded a segmentation error in the identification of the nuclei of 6.40 ± 1.07% (SFig.1E). We did not observe any substantial error additional to the nuclei segmentation error when we identified the Sox10^+^-GFP^+^ nuclei by the masking option (SFig.1E). Our analysis is a reliable tool for the fast quantification of cells in large image datasets. We calculated the percentage of cells belonging to the oligodendrocyte lineage over the total number of cells (DAPI^+^ nuclei) and detected the highest and lowest percentage in the corpus callosum and cortex at both P20 (65.87 ± 2.26% and 10.3 ± 0.62%) and P60 (67.95 ± 4.78% and 20.0 ± 5.13%. SFig.1D). This is in line with the previously described distribution of the OL lineage^50^ and the relatively myelination levels of the analyzed regions. The image analysis pipeline is available at the following link: https://github.com/Castelo-Branco-lab/Floriddia_et_al_2019.

### Clustering analysis

Cells were analyzed using the scater package^51^. Cells with less than 100 UMI counts and more than 15,000 were not considered for downstream analysis. Pool normalization was then used for normalization the expression matrix,^52^ feature selection was performed by calculating the coefficient of variation and using support vector regression to predict variance for a given gene expression level, all genes showing higher variance than expected were used for downstream analysis. The GeneFocus pipeline^18^ was used for gene filtering and Spearman correlation was used to build a KNN-graph after which Louvain clustering was applied. Cells that did not include any shared nearest neighbors were discarded from the clustering as outliers. Due to the limited number of cells present in some populations, we performed additional hierarchical clustering and set the threshold to separate the three main populations, oligodendrocyte progenitor cells (OPCs), committed oligodendrocyte progenitor cells (COPs), and mature oligodendrocytes (MOLs). The hierarchical and Louvain clusters were then combined, where the addition of the hierarchical clustering results led to the separation of COP cluster from the broader original Louvain clusters containing both OPCs and COPs. Code used for scRNAseq analysis is available at https://github.com/Castelo-Branco-lab/GeneFocus.

### Statistics

Statistics on the spatial distribution of the OL lineage populations was performed using 2-way ANOVA. For multiple comparison analysis, the Sidak’s correction was applied.

Differential gene expression analysis was performed using pairwise Wilcoxon rank sum tests using the stats package in R, on averaged expression per cluster. Significant genes were selected with a FDR adjusted p-value <0.01. The heatmap displays the most top 20 highest enriched genes as measured by z-score.

**Extended Data Fig. 1. Spatial distribution of OL populations in the juvenile and adult central nervous system.** (**A-B**) Schematics of coronal sections of the mouse brain (A) and spinal cord (B). Red squares highlight the systematically analyzed CNS regions. (**C**) Schematic overview and representative analyzed image output of the customized CellProfiler pipeline used in this study. (**D, F**) Percentage of the OL lineage cells (Sox10^+^ cells) calculated on the total number of nuclei shows the expected differential enrichment of the OL lineage in the analyzed regions. Data are presented as Mean ± SEM. n = 5-8 animals per condition. (**E**) Percentage of the segmentation error of the used pipeline. Data are presented as Mean ± SEM. n = 5-18 images per condition. (**G-H, K-R**) Confocal representative images of the distribution of OPC-COPs (Ptprz1^+^ OL lineage cells. G, K, O), MOL1 (Egr2^+^ OL lineage cells. H, L, P), MOL2 (Klk6^+^ OL lineage cells. M, Q), and MOL5/6 (Ptgds^+^ OL lineage cells. N, R) in the dorsal horn (grey matter) of the juvenile (P20) and adult (P60) spinal cord (G-H), cortex (K-N), and corpus callosum (O-R). Scale bar = 20 μm. (**I-J**) Quantification of the OPCs-COPs (I), MOL1 (J) distribution in the cortex, corpus callosum, and dorsal horn in juvenile (P20) and adulthood (P60). Percentage of the population is calculated on the total number of OL lineage cells (Sox10^+^ cells) in the analyzed region. Data are presented as Mean ± SEM. n = 6-9 animals per condition. Asterisks indicate a significant difference between conditions (*p ≤ 0.05, **p ≤ 0.01, ***p ≤ 0.001, ****p ≤ 0.0001, 2-way ANOVA with Sidak’s correction). Cx = cortex, CC = corpus callosum, SC = spinal cord, DC = dorsal columns, dCST = dorsal corticospinal tract.

**Extended Data Fig. 2. Louvain clustering analysis of the OL lineage cells.** (**A**) Violin plots depicting the expression of canonical markers for OPCs, MOL1, MOL2 and MOL5/6; (**B**) Heat-map of differential gene expression highlighting the enriched gene modules (right) characterizing the identified clusters (below) by Louvain clustering analysis. Distance matrix = Spearmann correlation.

**Extended Data Fig. 3. Pre- and postnatal OPCs equally contribute to the generation of MOL populations in the juvenile and adult central nervous system.** (**A-B**) Percentage of the fate mapped OL lineage cells (Sox10^+^-GFP^+^ cells) derived by OPCs labeled at E12.5 and P3-5 calculated on the total number of OL lineage cells (Sox10^+^ cells) at juvenile (A) and adulthood (B). (**C-F, H-N**) Confocal representative images show embryonically (TM E12.5)- and postnatally (TM P3-5)-derived OPCs-COPs (Ptprz1^+^-GFP^+^ OL lineage cells. C, H, K), MOL5/6 (Ptgds^+^-GFP^+^ OL lineage cells. D, L), MOL2 (Klk6^+^-GFP^+^ OL lineage cells. E, I, M), MOL1 (Egr2^+^-GFP^+^ OL lineage cells. F, J, N) in the adult (P60) cortex (C-F), corpus callosum (H-J), and dorsal horn of the spinal cord (K-N). Scale bar = 20 μm. (**G, O**) Percentages of the fate mapped OPCs-COPs (Ptprz1^+^-GFP^+^ OL lineage cells), MOL1 (Egr2-GFP^+^ OL lineage cells), MOL2 (Klk6^+^-GFP^+^ OL lineage cells), and MOL5/6 (Ptgds^+^-GFP^+^ OL lineage cells) populations are calculated on the total number of fate mapped OL lineage cells (Sox10^+^-GFP^+^ cells) in the juvenile and adult cortex (G) and spinal cord (O). (**P-Q**) Percentages of the embryonically (TM E12.5)- and postnatally (TM P3-5)-derived MOL2 (Klk6^+^-GFP^+^ OL lineage cells. P), and MOL5/6 (Ptgds^+^-GFP^+^ OL lineage cells. (Q) populations are calculated on the total number of fate mapped OL lineage cells (Sox10^+^-GFP^+^ cells) in the spinal cord white and grey matter. Data are presented as Mean ± SEM. n = 3-5 animals per condition. Asterisks indicate a significant difference between conditions (*p ≤ 0.05, **p ≤ 0.01, ***p ≤ 0.001, ***p ≤ 0.0001, 2-way ANOVA with Sidak’s correction). TM = tamoxifen, GFP = green fluorescent protein, GM = grey matter, WM = white matter.

**Extended Data Fig. 4. MOL2 population expands away form the injury site but maintain its spatial preference following spinal cord injury.** (**A**) Confocal representative images of the dorsal spinal cord region 10-5 mm rostral to the injury site showing the maintained MOL2 (Klk6^+^ OL lineage cells) and MOL5/6 (Ptgds^+^ OL lineage cells) spatial preference 5 months post-injury. White dashed lines highlight the dorsal funiculi. Scale bar = 100 μm. (**B-E**) Percentages of the MOL2 (Klk6^+^ OL lineage cells) and MOL5/6 (Ptgds^+^ OL lineage cells) populations are calculated on the total number of OL lineage cells (Sox10^+^ cells) in the dorsal columns (B-C) and dorsal corticospinal tract (D-E) 5-10mm rostral and caudal to the injury site. Data are presented as Mean ± SEM. n = 5-7 animals per condition. Asterisks indicate a significant difference between conditions (*p ≤ 0.05, **p ≤ 0.01, ***p≤ 0.001, 2-way ANOVA with Sidak’s correction). mpi = months post-injury.

**Extended Data Table 1. Quantification of the OL subpopulations contribution to the OL lineage.** Percentage of the OPC-COPs (Ptprz1^+^ OL lineage cells), MOL1 (Egr2^+^ OL lineage cells), MOL2 (Klk6^+^ OL lineage cells), and MOL5/6 (Ptgds^+^ OL lineage cells) contribution to the OL lineage (Sox10^+^ cells) in the regions of interest in juvenile (P20) and adulthood (P60). We imaged 0.3, 0.13, and 0.4 mm2 of the sensorimotor cortex, corpus callosum, and dorsal spinal cord per tissue section, respectively. Minimum three sections per animal were analyzed. Data are presented as Mean ± SEM. n = 4-9 animals per condition.

**Extended Data Table 2. Average number of the quantified cells per analyzed section.** Average number of the OL lineage cells (Sox10^+^) and OPC-COPs (Ptprz1^+^ OL lineage cells), MOL1 (Egr2^+^ OL lineage cells), MOL2 (Klk6^+^ OL lineage cells), and MOL5/6 (Ptgds^+^ OL lineage cells) counted per region of interest in juvenile (P20) and adulthood (P60) per section. We imaged 0.3, 0.13, and 0.4 mm2 of the sensorimotor cortex, corpus callosum, and dorsal spinal cord per tissue section, respectively. Mimimum three sections per animal were analyzed. Data are presented as Mean ± SEM. n = 4-9 animals per condition.

**Extended Data Table 3. Counting of analyzed cells by scRNAseq.**

**Extended Data Table 4. Quantification of the OL subpopulations contribution to the OL lineage.** Percentage of the MOL2 (Klk6^+^ OL lineage cells) and MOL5/6 (Ptgds^+^ OL lineage cells) contribution to the OL lineage (Sox10^+^ cells) in the spinal cord regions in juvenile (P20) and adulthood (P60). Minimum three sections per animal were analyzed. Data are presented as Mean ± SEM. n = 5-7 animals per condition.

## Footnotes

1 The following secure token has been created to allow review of record GSE128525 while it remains in private status: kfizogqqrhwpnwn

