## Supplementary Figures and Tables for "Distinct oligodendrocyte populations have spatial preference and injury-specific responses"

Floriddia et al., Extended Data Fig.1

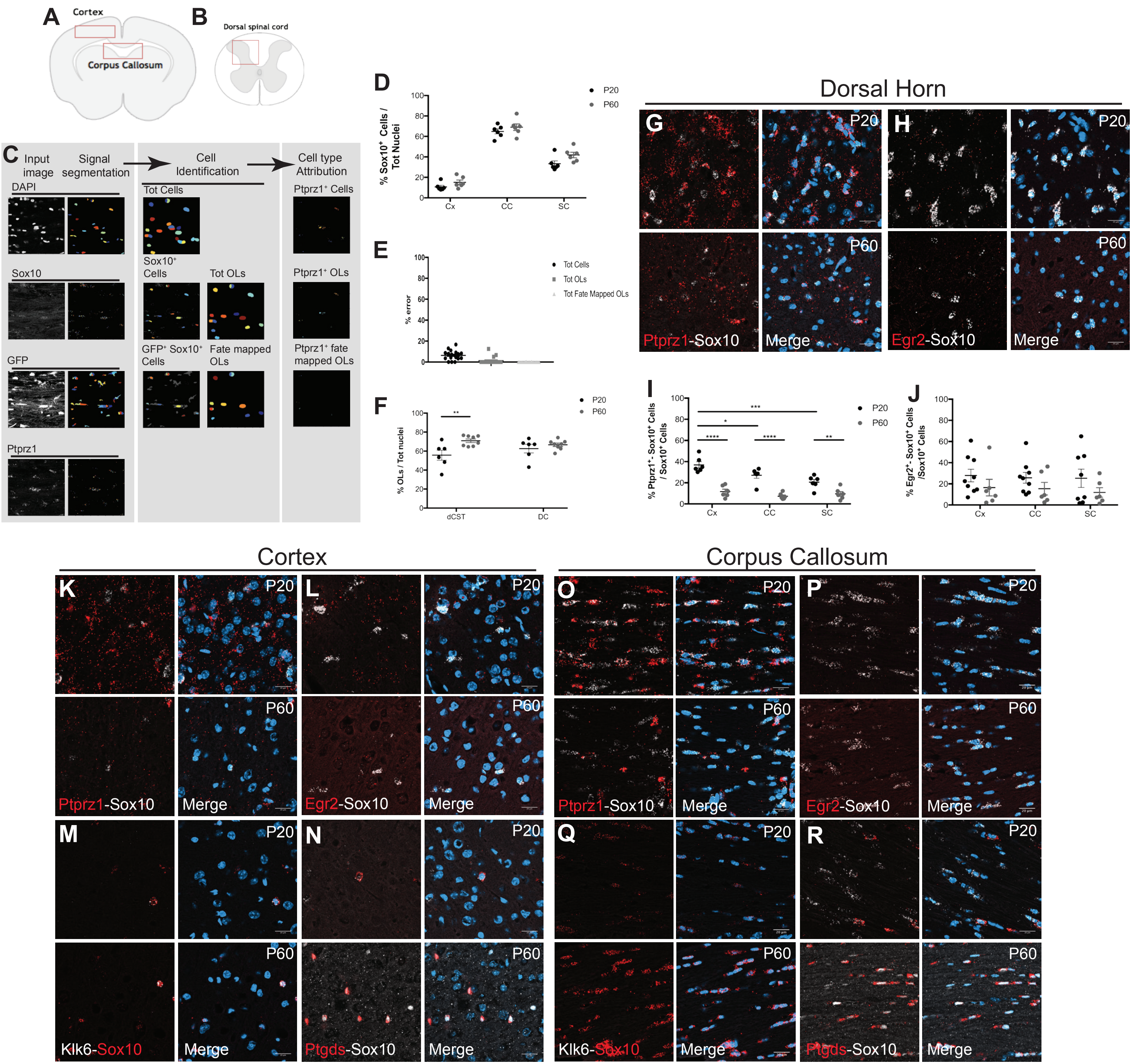

Floriddia et al,  Extended Data Fig. 2

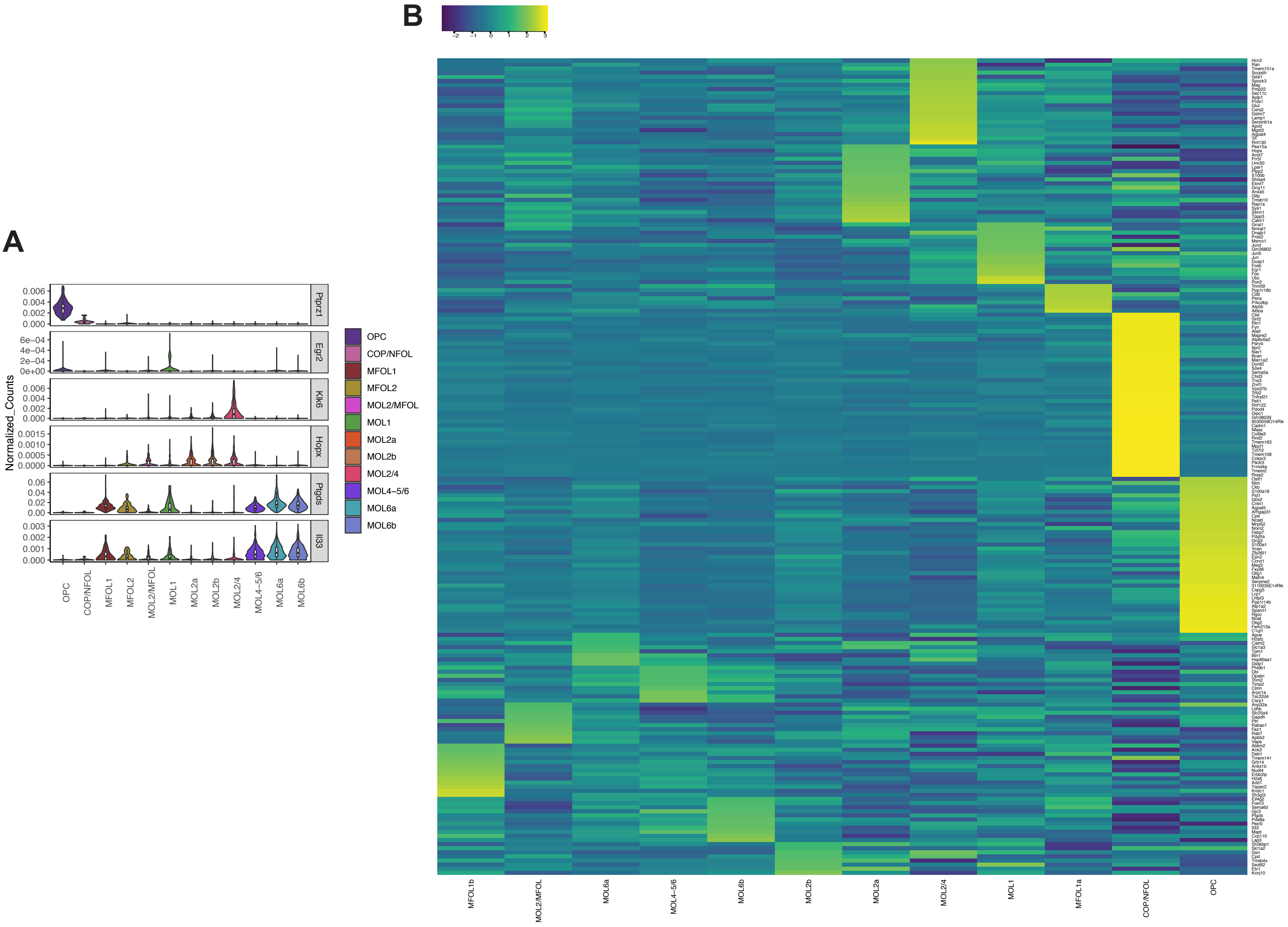

Floriddia et al,  Extended Data Fig. 3

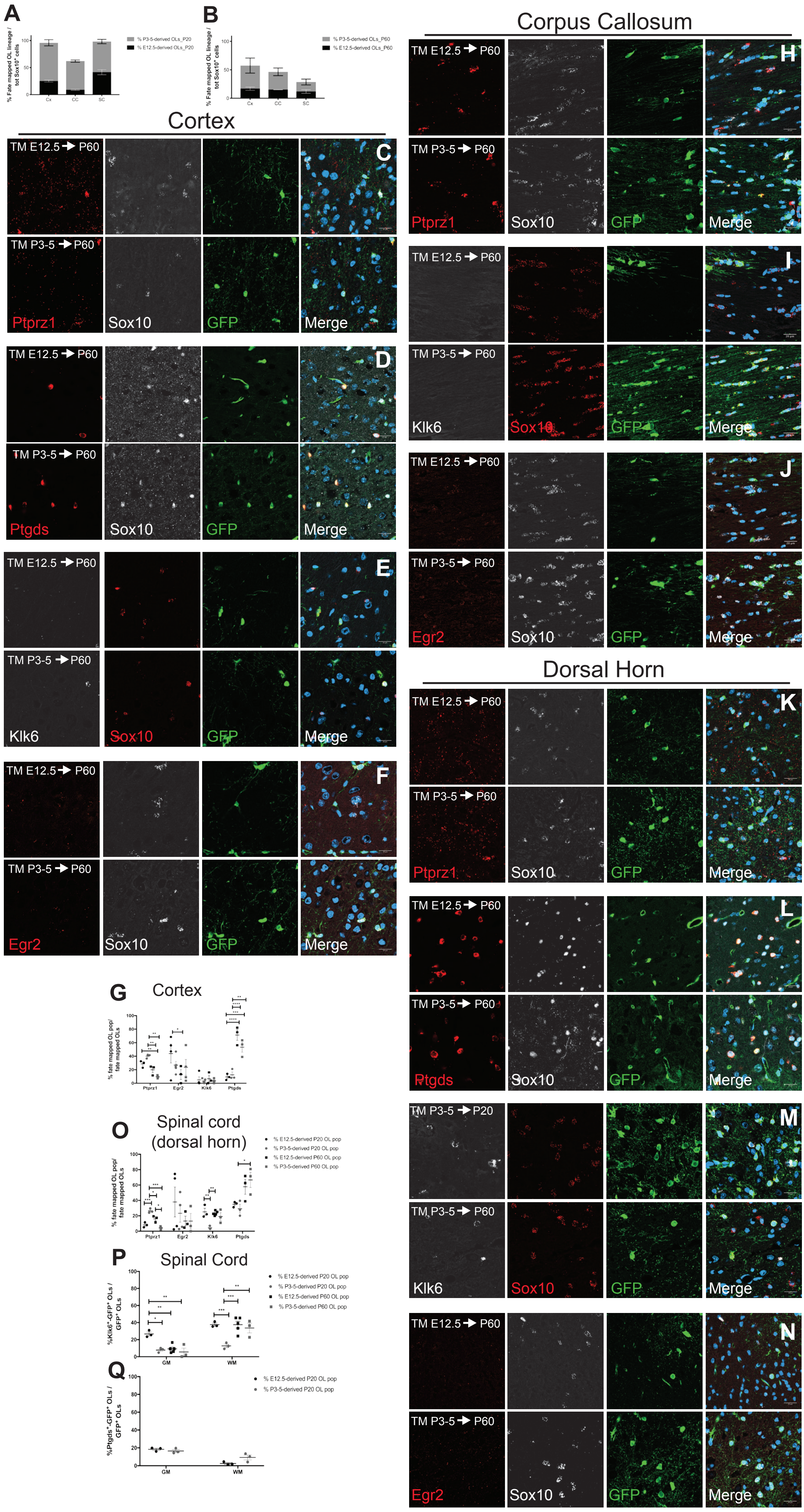

Floriddia et al., Extended Data Fig. 4

10-5 mm Rostral to lesion

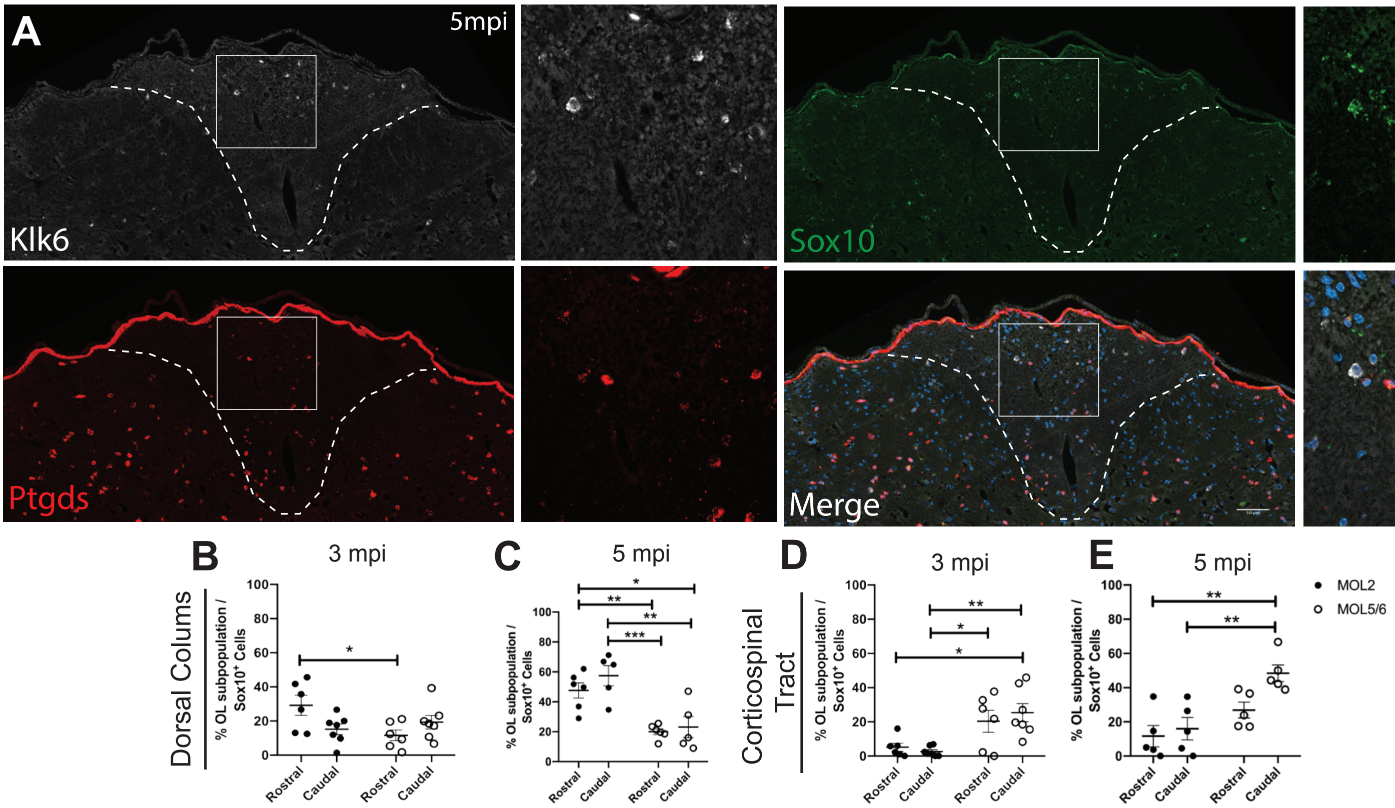

| CNS region | Age | OL lineage subpopulation | Contribution to the lineage (% ± SEM) |
|---|---|---|---|
| Somatosensory cortex | P20 | OPCs/COPs | 36.91 ± 2.98 |
| Corpus callosum | P20 | OPCs/COPs | 27.14 ± 2.85 |
| Dorsal horn | P20 | OPCs/COPs | 20.61 ± 2.48 |
| Somatosensory cortex | P60 | OPCs/COPs | 11.73 ± 2.23 |
| Corpus callosum | P60 | OPCs/COPs | 7.25 ± 1.19 |
| Dorsal horn | P60 | OPCs/COPs | 9.55 ± 2.16 |
| Somatosensory cortex | P20 | MOL1 | 27.83 ± 5.93 |
| Corpus callosum | P20 | MOL1 | 25.71 ± 5.05 |
| Dorsal horn | P20 | MOL1 | 25.26 ± 8.14 |
| Somatosensory cortex | P60 | MOL1 | 16.46 ± 7.87 |
| Corpus callosum | P60 | MOL1 | 15.44 ± 5.97 |
| Dorsal horn | P60 | MOL1 | 11.89 ± 4.51 |
| Somatosensory cortex | P20 | MOL2 | 3.49 ± 0.96 |
| Corpus callosum | P20 | MOL2 | 0.23 ± 0.12 |
| Dorsal horn | P20 | MOL2 | 15.34 ± 2.36 |
| Somatosensory cortex | P60 | MOL2 | 3.24 ± 0.97 |
| Corpus callosum | P60 | MOL2 | 0.39 ± 0.18 |
| Dorsal horn | P60 | MOL2 | 16.27 ± 1.37 |
| Somatosensory cortex | P20 | MOL5/6 | 10.24 ± 1.84 |
| Corpus callosum | P20 | MOL5/6 | 9.39 ± 2.08 |
| Dorsal horn | P20 | MOL5/6 | 24.71 ± 3.08 |
| Somatosensory cortex | P60 | MOL5/6 | 54.43 ± 3.51 |
| Corpus callosum | P60 | MOL5/6 | 63.77 ± 6.31 |
| Dorsal horn | P60 | MOL5/6 | 51.37 ± 3.14 |

| CNS region | Age | Cell type | Number of cells per section (avg. ± SEM) |
|---|---|---|---|
| Somatosensory cortex | P20 | OL lineage | 70 ± 5 |
| Corpus callosum | P20 | OL lineage | 151 ± 15 |
| Dorsal horn | P20 | OL lineage | 198 ± 21 |
| Somatosensory cortex | P60 | OL lineage | 61 ± 7 |
| Corpus callosum | P60 | OL lineage | 164 ± 12 |
| Dorsal horn | P60 | OL lineage | 164 ± 23 |
| Somatosensory cortex | P20 | OPCs/COPs | 19 ± 2 |
| Corpus callosum | P20 | OPCs/COPs | 39 ± 7 |
| Dorsal horn | P20 | OPCs/COPs | 35 ± 7 |
| Somatosensory cortex | P60 | OPCs/COPs | 12 ± 2 |
| Corpus callosum | P60 | OPCs/COPs | 14 ± 4 |
| Dorsal horn | P60 | OPCs/COPs | 23 ± 6 |
| Somatosensory cortex | P20 | MOL1 | 15 ± 5 |
| Corpus callosum | P20 | MOL1 | 40 ± 12 |
| Dorsal horn | P20 | MOL1 | 65 ± 21 |
| Somatosensory cortex | P60 | MOL1 | 10 ± 5 |
| Corpus callosum | P60 | MOL1 | 27 ± 10 |
| Dorsal horn | P60 | MOL1 | 16 ± 6 |
| Somatosensory cortex | P20 | MOL2 | 2 ± 1 |
| Corpus callosum | P20 | MOL2 | 1 ± 0 |
| Dorsal horn | P20 | MOL2 | 31 ± 7 |
| Somatosensory cortex | P60 | MOL2 | 2 ± 1 |
| Corpus callosum | P60 | MOL2 | 1 ± 0 |
| Dorsal horn | P60 | MOL2 | 29 ± 6 |
| Somatosensory cortex | P20 | MOL5/6 | 5 ± 1 |
| Corpus callosum | P20 | MOL5/6 | 10 ± 3 |
| Dorsal horn | P20 | MOL5/6 | 54 ± 22 |
| Somatosensory cortex | P60 | MOL5/6 | 37 ± 5 |
| Corpus callosum | P60 | MOL5/6 | 131 ± 5 |
| Dorsal horn | P60 | MOL5/6 | 93 ± 20 |

**Table 1**

| | OPC | COP/NFOL | MFOL1 | MFOL2 | MOL2/MFOL | MOL1 | MOL2a | MOL2b | MOL2/4 | MOL4-5/6 | MOL6a | MOL6b | Total |
|---|---|---|---|---|---|---|---|---|---|---|---|---|---|
| **TdTom+ OL lineage cells** | 48 | 11 | 175 | 47 | 170 | 111 | 163 | 270 | 60 | 252 | 300 | 294 | 1901 |
| **GFP+ OL lineage cells** | 35 | 8 | 99 | 19 | 61 | 50 | 67 | 165 | 63 | 98 | 151 | 136 | 952 |
| **Tot analyzed OL lineage cells** | 83 | 19 | 274 | 66 | 231 | 161 | 230 | 435 | 123 | 350 | 451 | 430 | 2853 |

| Spinal cord region | Age | OL lineage subpopulation | Contribution to the lineage (% ± SEM) |
|---|---|---|---|
| Grey matter | P20 | MOL2 | 15.26 ± 1.87 |
| White matter | P20 | MOL2 | 25.13 ± 2.31 |
| Grey matter | P60 | MOL2 | 7.16 ± 1.15 |
| White matter | P60 | MOL2 | 24.83 ± 1.92 |
| Grey matter | P20 | MOL5/6 | 30.07 ± 3.19 |
| White matter | P20 | MOL5/6 | 14.31 ± 2.04 |
| Grey matter | P60 | MOL5/6 | 51.37 ± 3.14 |
| White matter | P60 | MOL5/6 | 11.97 ± 2.78 |
| Dorsal corticospinal tract | P20 | OL lineage | 55.73 ± 5.29 |
| Dorsal columns | P20 | OL lineage | 62.65 ± 4.5 |
| Dorsal corticospinal tract | P60 | OL lineage | 70.91 ± 1.9 |
| Dorsal columns | P60 | OL lineage | 66.64 ± 1.8 |
| Dorsal corticospinal tract | P20 | MOL2 | 5.33 ± 1.35 |
| Dorsal columns | P20 | MOL2 | 19.32 ± 3.18 |
| Dorsal corticospinal tract | P60 | MOL2 | 5.25 ± 0.8 |
| Dorsal columns | P60 | MOL2 | 24.26 ± 1.83 |
| Dorsal corticospinal tract | P20 | MOL5/6 | 19.69 ± 6.67 |
| Dorsal columns | P60 | MOL5/6 | 12.88 ± 4.82 |
| Dorsal corticospinal tract | P60 | MOL5/6 | 50.91 ± 8.37 |
| Dorsal columns | P60 | MOL5/6 | 18.78 ± 1.97 |
